# Direct measurement of PIP_2_ densities in biological membranes using a peptide-based sensor

**DOI:** 10.1101/2024.09.11.612554

**Authors:** Vinay K. Menon, Joy Wu, Alex J. Alonzo, Kaitlyn A. Rogers, Kevin L. Scrudders, Suriya Selvarajan, Andrew Walke, Rajasree Kundu, Ankona Datta, Shalini T. Low-Nam

## Abstract

Organization and composition of the plasma membrane are important modulators of many cellular programs. Phosphatidylinositol phosphate (PIP) lipids are low abundance membrane constituents with different arrangements of phosphate groups around an inositol head group that regulate a large number of signaling pathways. Many strategies have been developed to detect and track PIP species to monitor their clustering, mobility, and interaction with binding partners. We implement a peptide-based ratiometric sensor for the detection of PI(4,5)P_2_ lipids in reconstituted membrane systems that permit absolute quantification of PI(4,5)P_2_ densities down to physiological levels. The sensor is membrane permeable and easily applicable to measurements in living cells. Application of calibrated sensors to cells expressing common mutations in the small GTPase, Ras, showed a reshaping of surface PI(4,5)P_2_ levels and distributions in a mutation-specific manner. The rapid implementation of this quantitative sensing strategy to cellular studies of cellular signaling, membrane organization and dynamics should be broadly applicable.

## Introduction

Despite very low abundance in the plasma membrane, phosphatidylinositol phosphate (PIP) lipids, generate specific signaling outcomes ranging from actin reorganization to immune signaling ((1–5)). PIP lipid species, defined by unique arrangements of phosphate groups around an inositol head group, predominate the cytoplasmic leaflet of the plasma membrane and comprise approximately 2 mole percent of the total lipid content, based on lipidomic data (6). Phosphatidylinositol-4,5-bisphosphate (PI(4,5)P_2_) is the most abundant PIP species and plays a key role in the activation of the guanine nucleotide exchange factor (GEF) son of sevenless (SOS) through pleckstrin homology (PH) domain-mediated release of autoinhibition ((7, 8)). Active SOS primes activation of the small GTPase, Ras, and drives mitogenic signaling. Ras mutations are common drivers in many cancers which has led to extensive efforts to block oncogenic Ras signaling. The recent development of inhibitors specific for Ras G12 allele-specific mutations have been a major advance in lung cancer therapy ((9–11)).

As a substrate, PI(4,5)P_2_ is cleaved into second messengers inositol triphosphate (IP_3_) and diacylglycerol (DAG) by phospholipase C, and is converted to PI(3,4,5)P_3_ by PI3K ((12–15)). PI(3,4,5)P_3_ activates downstream signaling pathways involved in cell growth and survival and the polarized redistribution of PI(4,5)P_2_ and PI(3,4,5)P_3_ is a feature of polarized cell migration (refs). Thus, PI(4,5)P_2_ is required as an activator for certain membrane-associated programs but, in parallel, it is depleted by competing processes. Adding to the complex dynamics at the membrane, Ras itself interacts with PI(4,5)P_2_ and recruits PI3K to the surface (15, 16). Given the dependence of many signaling programs on local PI(4,5)P_2_ abundance, detection of the rare lipid has been the subject of great interest.

Heterogeneity in PI(4,5)P_2_ distribution has been identified in polarized cell migration and endocytosis whose collective processes span the nanometer-to-micron length scale and seconds to minutes time scale (1–5, 17). Measurements of PI(4,5)P_2_ mobility and dynamics have mainly been based on fluorescently-tagged binding domains or antibody-based detection. The PH domain from phospholipase C delta 1 (PH-PLCδ1) is commonly used to track PI(4,5)P_2_ mobility and has shown both rapid and slow diffusive states ((18)). Restricted diffusion has been implicated in PI(4,5)P_2_ signaling outcomes and emphasizes how local enhancement in PI(4,5)P_2_ may overcome the overall low abundance (19). However, recent work using the C-terminal domain of Tubby showed that PH-PLCδ1 exhibits slower diffusion of PI(4,5)P_2_ by an unknown interaction. Other challenges in detecting PI(4,5)P_2_ using these domain-based biosensors include their relatively large sizes and the requirement for microinjection, electroporation, or transfection into cells for visualization (20). Thus, characterization of PI(4,5)P_2_ distributions and dynamics across a population of cells may be impaired by the technical challenges in using these biosensors.

Datta and colleagues recently introduced a cell penetrating, ratiometric, peptide-based PI(4,5)P_2_ sensor, DAN13aa (20). The short, cationic sequence, derived from the actin-binding protein gelsolin, penetrates bilayer membranes and is selective for PI(4,5)P_2_ (20). To calibrate the DAN13aa biosensor, we developed an *in vitro* reconstitution-based approached using supported lipid bilayers (SLBs) across a range of PI(4,5)P_2_ densities, including the low levels that approach predicted physiological densities (6). Fast binding dynamics and a linear responsiveness of the sensor permitted characterization of PI(4,5)P_2_ distributions in living cells expressing point mutations of KRas4B in an isogenic background. Focusing on KRas4B mutations with impaired GTPase activity that exacerbate PI3K activation (21), an unexpected heterogeneity emerged. Both levels of PI(4,5)P_2_ and organization at the membrane were altered at steady state. Only G12D mutant cells had diminished PI(4,5)P_2_ levels consistent with largescale conversion to PI(3,4,5)P_3_. We propose that an increase in spatial heterogeneity of phosphoinositide distribution is a hallmark of oncogenesis.

## Results

### Stacked Supported Lipid Bilayers (SSLB) Prevents DAN13aa Substrate Sticking

During its development, the DAN13aa sensor was shown to have selectivity for PI(4,5)P_2_ over PI(4)P and a micromolar affinity (20). The sensor could be ideally suited to PI(4,5)P_2_ detection in cellular membranes with a strong sensitivity to the physiologically low levels of the lipid in the plasma membrane. To monitor sensor responsiveness at PI(4,5)P_2_ densities below 5 mole percent, we developed an *in vitro* reconstituted platform based on supported lipid bilayers (SLBs) and implemented surface-selective imaging in a total internal reflection (TIR) fluorescence configuration. Measurements were made with the sensor in solution to establish a steady state of exchange between the membrane and the bulk. The 13-mer peptide is conjugated to the solvatochromic fluorophore, 2-dimethylamino-6-acyl-naphthalene (DAN) at the N-terminal cysteine residue. Dual wavelength detection was used, following 405 nm excitation, to monitor the shift from green emission in an aqueous environment (unbound, λ_em_max_ = 520 nm) to blue emission in a hydrophobic environment (bound, λ_em_max_ = 450 nm; Fig. 1A (20)). Data were represented as bound fraction based on the ratio of bound:unbound signal. In initial measurements, PI(4,5)P_2_-containing SLBs were formed by vesicle fusion on a negatively-charged solid glass support. Consistent with a strong membrane permeability, the positively-charged DAN13aa sensor passed easily through SLBs and stuck non-specifically to the underlying glass substrate, even in the absence of PI(4,5)P_2_ (Fig 1B, top; Fig. S1).

**Figure 1.**
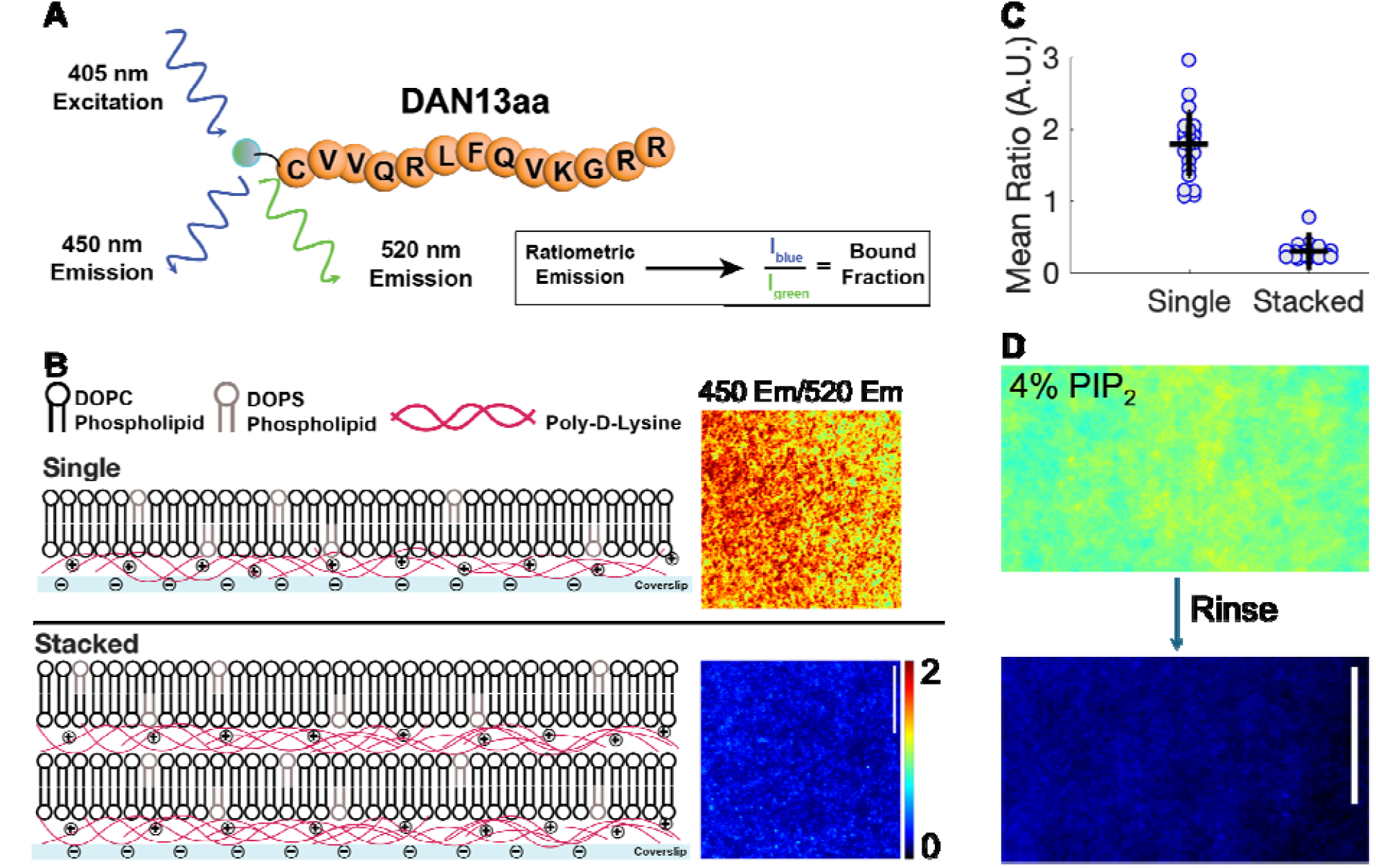
DAN13aa sensor sensitively detects PI(4,5)P2 in stacked supported lipid bilayers. A. The DAN13aa sensor is detected ratiometrically using 405 nm excitation and emission in 450 nm and 520 nm. B. Schematics of single and stacked SLBs on poly-lysine-coated glass substrates. In each case, the ratio of emissions is shown for a representative field of view. C. Quantification of mean ratio in single and stacked SLB configurations for at least 2 substrates and multiple fields of view. D. Ratiometric signal for a 4% PIP2-containing top bilayer in a stacked configuration before (top) and after (bottom) rinsing with buffer. All scale bars are 10 μm.

Using a previously-described strategy, we introduce passivated, stacked SLBs as a reconstituted membrane platform (22). The stacked SLBs permit DAN13aa sensing without non-specific sensor accumulation. The stacked SLBs consisted of two supported lipid bilayers deposited alternately on top of layers of poly-D-lysine (Fig. 1B). The stacked SLB (SSLB) platform ensured that the cationic poly-lysine repelled the positively charged DAN13aa sensor and resulted in a dramatic reduction in background in the absence of PI(4,5)P_2_ in the bilayers (Fig. 1B-C). The sensor has a negligible interaction with with DOPC and DOPS (20). Subsequent *in vitro* experiments were conducted using SSLBs.

Mobility of each of the bilayers in the SSLBs was confirmed by fluorescence recovery after photobleaching (FRAP; Fig. S2). Single lipid mobility in the upper bilayer was also verified by doping in 1 mole percent of fluorescently-tagged PI(4,5)P_2_. Following photobleaching to achieve single molecule densities, individual lipids were easily detected diffusing in the bilayer (Fig. S3). Notably, the second poly-D-lysine layer was found to be mobile, presumably due to the fluidity of the underlying SLB. Using fluorescently-tagged PI(4,5)P_2_ lipids and single particle tracking, we found that the individual phosphoinositide lipids were mobile in the upper SLB. Thus, the SSLB configuration also overcomes the slowed mobility of PI(4,5)P_2_ found in single SLBs proposed to be the consequence of lipid flipping to the lower leaflet and strong hydrogen bonding with the glass substrate (23).

The ideal application of the DAN13aa sensor to PI(4,5)P_2_ dynamics would minimize perturbation to normal processes at the membrane. Competition for PI(4,5)P_2_ binding or long interaction lifetimes could alter normal signaling processes. We verified rapid exchange of DAN13aa sensor on SSLBs with 4 mole percent PI(4,5)P_2_. The measured bound fraction rapidly returned to background levels following a rinse with buffer, suggesting fast binding kinetics (Fig. 1D).

### SSLBs Enable Precise Calibration of the DAN13aa PI(4,5)P2 Biosensor

PI(4,5)P2 densities in cellular membranes have been estimated to be approximately 20,000 molecules/µm^2^ and 8 × 10^6^ - 6.8 × 10^7^ molecules in the inner leaflet of mammalian cells (24–28). Calibration of the DAN13aa biosensor in SSLBs ranging in PI(4,5)P_2_ densities from 0-10 mole percent spanned the pertinent range expected in the plasma membrane of cells (Fig. 2A). We confirmed the phosphoinositide lipid composition in the small unilamellar vesicles used in the formation of SSLBs by a phosphate quantification assay. Digestion of lipid mixtures by perchloric acid results in free phosphates that are quantified by a colorimetric reaction with a manganese moiety (29). Using vesicles that contain 0, 1, and 4 mole percent porcine brain PI(4,5)P_2_, we showed that the phosphate levels were tightly confined to the expected amounts of the phosphoinositide (Fig. S4).

**Figure 2.**
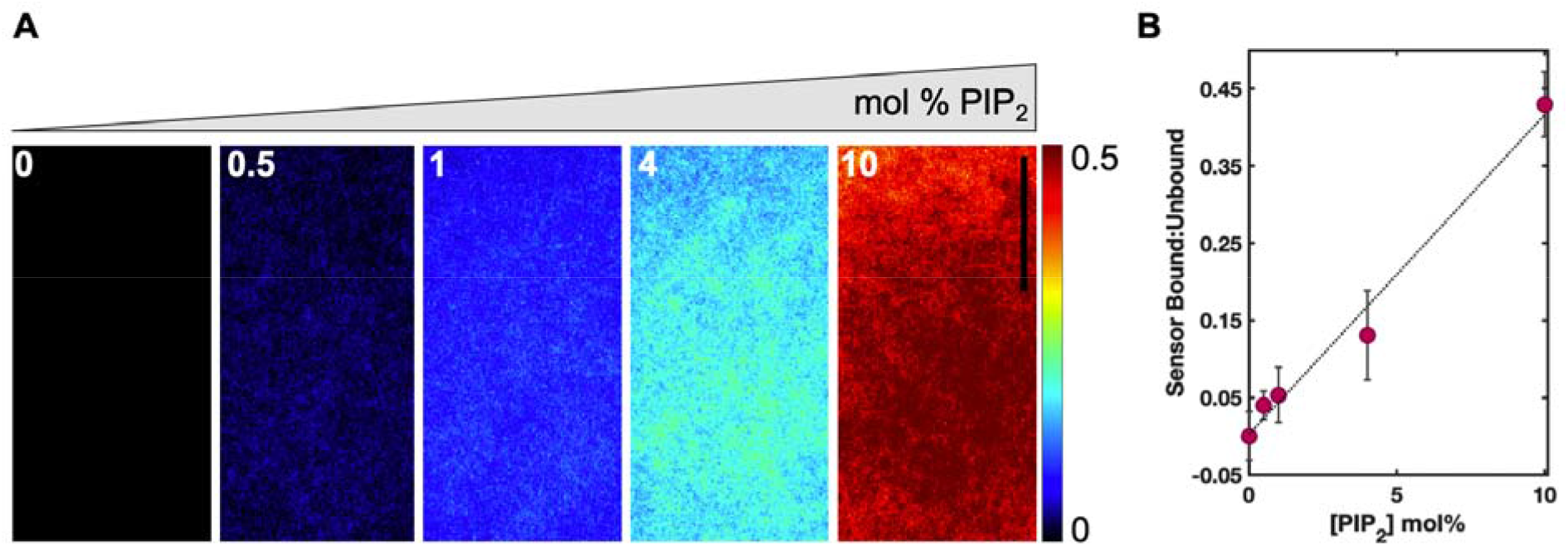
Calibration of DAN13aa in stacked supported lipid bilayers (SSLBs). A. SLBs were constructed. All samples contained 1 mol% DOPS/99 mol% DOPC in the bottom bilayer and 1 mol% DOPS/99 mol% DOPC; 0.5 mol% PI(4,5)P_2_/99.5 mol% DOPC; 1 mol% PI(4,5)P_2_/99 mol% DOPC; 4 mol% PI(4,5)P_2_96 mol% DOPC; or 10 mol% PI(4,5)P_2_/90 mol% DOPC in the top bilayer, as indicated. Ratiometric signal from 405 nm excitation and emission collected at 405 or 488 nm is shown. B. The ratiometric intensity of 405-405 to 405-488, corresponding to the bound fraction of DAN13aa, was plotted as a function of PI(4,5)P_2_ mol% in the top bilayer and fit with a linear regression (All scale bars, 10 µm).

There was a very strong linear correlation (R^2^ = 0.984) between the PI(4,5)P_2_ content in the stacked bilayer and the corresponding ratiometric signal intensity from the DAN13aa biosensor (Fig. 2B). A typical challenge in designing phosphoinositide sensors is the high degree of structural similarity between the lipids. Given the role Ras in activating PI3K to convert PI(4,5)P_2_ into PI(3,4,5)P_3_, we measured the ability of DAN13aa to distinguish between these phosphoinositides. There was some promiscuity in sensor detection of PIP(3,4,5)_3_-containing SSLBs. In stacked SLBs each with 4 mole percent of phosphoinositide, the DAN13aa sensor detects PI(4,5)P_2_ with 2-3-fold selectivity over PIP(3,4,5)_3_ (Fig. S6). Thus, in cells, the application of DAN13aa to PI(4,5)P_2_ density is predominated by the dually-phosphorylated lipid but may measure the presence of PI(4,5)P_3,_ perhaps, in particular, detecting the local conversion to the latter by PI3K. Further application of the sensor to living cells is promoted by a negligible difference in sensor sensitivity at room temperature as compared to 37°C (Fig. S5).

### Application of the DAN13aa sensor PI(4,5)P2 levels and distributions in living cells

For experiments in cells, we used the isogenic Ras cell lines established in the mouse embryonic fibroblast (MEF) background (21). The cells are NRas and HRas null and include wild-type (WT) KRas4B or common clinical Ras mutants. A KRas4B null variant is dependent on a BRaf V600E mutation for growth. These cells provide a common background to measure the distribution of PI(4,5)P_2_ lipids in the plasma membrane under modulation by oncogenic Ras. Initial characterization of the sensor in cells was carried out using the WT cells as a model. Consistent with the high degree of membrane permeability designed into the cationic DAN13aa peptide, sensor uptake, measured by flow cytometry, was uniform across the cell population (Fig. 3A). Thus, we were able to measure PI(4,5)P_2_ features across large numbers of cells and without the need for expression or introduction of an exogenous biosensor. Toxicity of the sensor was determined using an MTT assay. Cells maintained a high level of viability even after 24 hours of exposure to the DAN13aa peptide.

**Figure 3.**
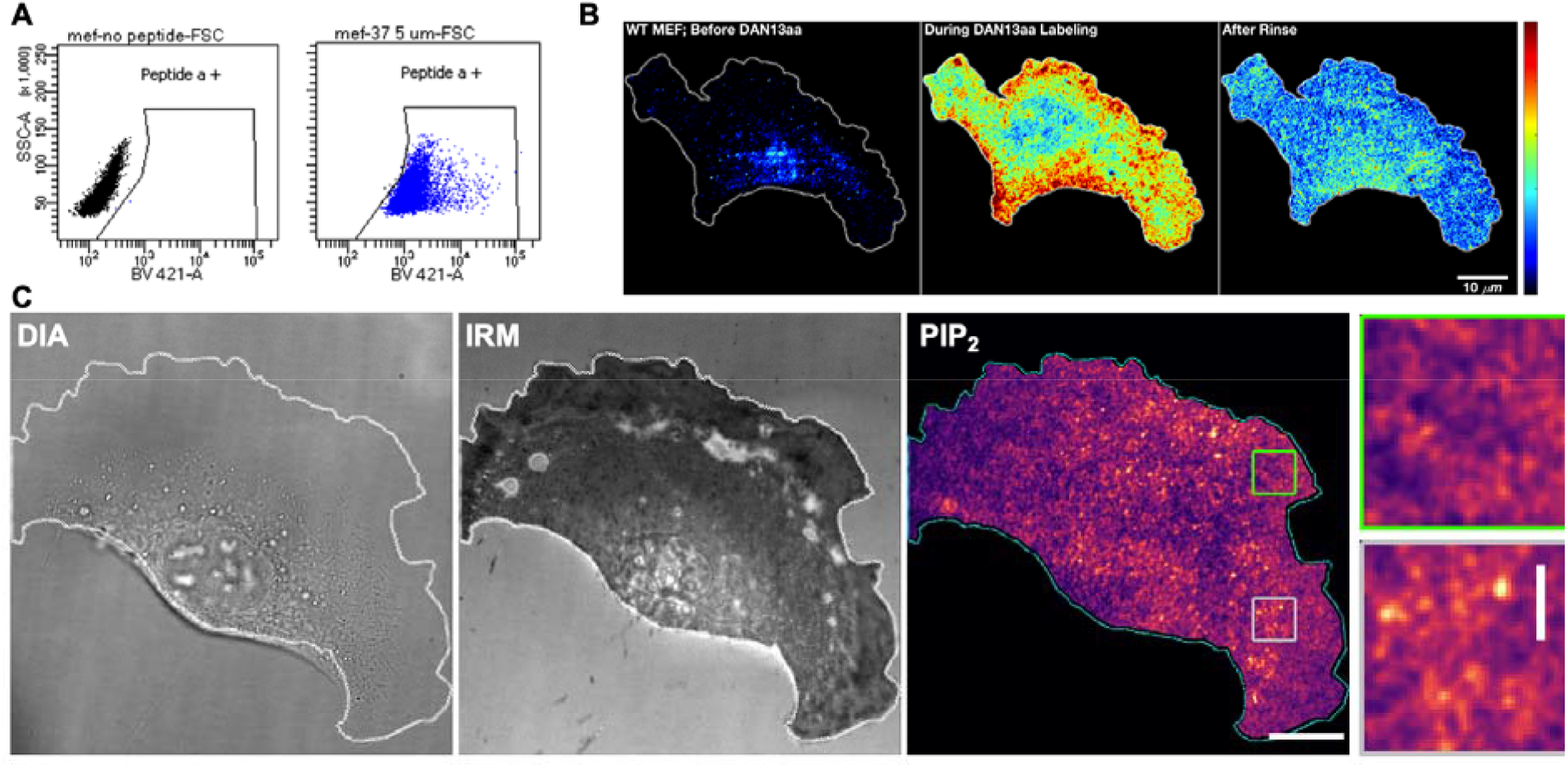
Labeling of cells with DAN13aa sensor. A. Flow cytometry of WT MEF cells without (left) and with (right) DAN13aa sensor. B. Labeling of WT MEF cells before (left) and during (middle) DAN13aa labeling. Following a rinse with buffer the steady state DAN13aa ratiometric signal was monitored (right). C. WT MEF cells showing brightfield image, adherent membrane imaged in interference contrast microscope (IRM) and PI(4,5)P_2_ density following adjustment of ratiometric signal using calibration curve. Regions of interest are enlarged to the right.

Labeling of cells was also visualized using ratiometric imaging in TIRF to detect PI(4,5)P_2_ lipids at the adherent membrane. A low level of ratiometric signal was observed prior to exposure to the DAN13aa sensor (Fig. 3B). The sensor was rapidly taken up by cells and reached a high level of signal. Following a short incubation and a rinse with buffer, cells retained an equilibrium amount of sensor and showed a measurable ratiometric signal (Fig. 3B-3C). Using the calibration from the SSLBs, the measured PI(4,5)P_2_ levels were converted to mole percentages. Overall, PI(4,5)P_2_ distributions in WT cells were uniform but some clustered lipids with local concentrations up to almost 1 mole percent were observed (Fig. 3C). Small clusters of PI(4,5)P_2_ lipids are consistent with previous observations (30, 31). We confirmed the distribution of PI(4,5)P_2_ lipids using immunofluorescent detection, which also showed some local clustering but a spread across the membrane (Fig. S7).

### PI(4,5)P2 densities in MEFs expressing Ras mutants

To extend the cellular observations to MEFs expressing Ras mutants, we selected the G12 and G13 mutations to focus on the mechanism of slowed Ras inactivation via interference of GAP binding. To avoid toxicity from extended exposure to 405 nm light, PI(4,5)P_2_ levels were monitored under steady state conditions. Observations of single cells of WT, G12D, G12V, and G13D genotypes, showed that the mean PI(4,5)P_2_ levels covered a narrow range, but were different for each genotype. However, when compared to WT cells, the mutant cells appeared to have greater heterogeneity in PI(4,5)P_2_ densities. This observation was inconsistent with a simple explanation that hyperactive Ras primes conversion of PI(4,5)P_2_ to PI(3,4,5)P_3_ by PI3K to promote leading edge protrusivity. Cells were observed under tonic signaling conditions which may have obscured some excitable activities that have been observed in Ras, PI3K, actin networks (32).

To characterize spatial patterning of PI(4,5)P_2_ distributions in the Ras mutant cells, we applied texture analysis using the gray-level co-occurrence matrix (GLCM). In this approach, features of neighboring pixels are extracted to assess the signal heterogeneity. GLCM analysis extracts the distribution of co-occurring pixel grayscale values across an image to show patterns and topography within an image (33, 34). When the mean PI(4,5)P_2_ intensities was plotted against the GLCM values, the genotypes were found to cluster into characteristic regions of the resulting density-spatial heterogeneity space (Fig. 4). G13D cells were found to have the highest degree of heterogeneity and WT cells had the most homogeneous PI(4,5)P_2_ distributions. The G12D and G12V cells had similar mean PI(4,5)P_2_ levels, but the latter had a slightly higher degree of heterogeneity in the phosphoinositide distribution. Segregation of the genotypes into regions of the information space suggests characteristic PI(4,5)P_2_ features may be correlated with specific cancers.

**Figure 4.**
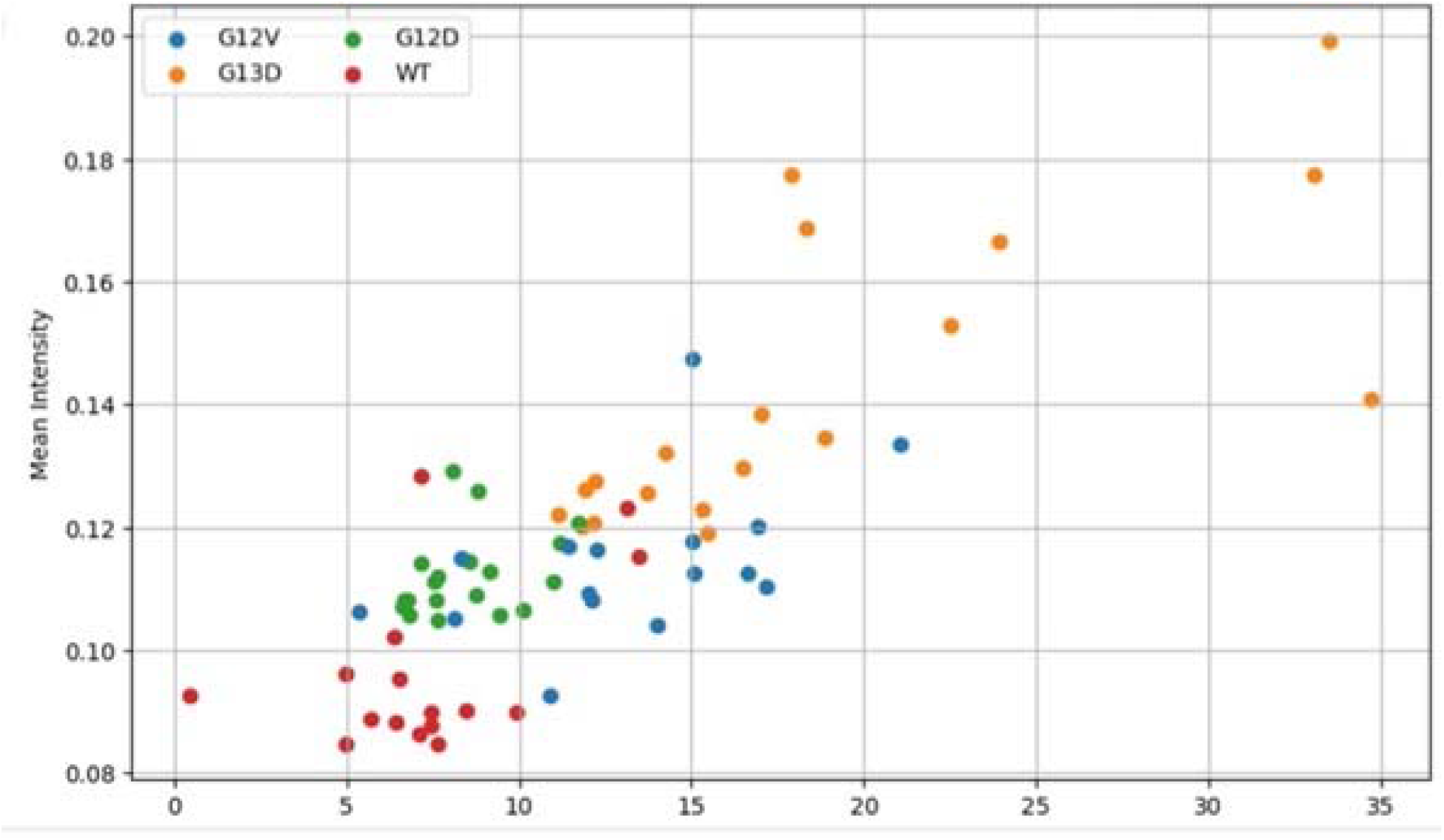
Clustering of PI(4,5)P_2_ densities in single Ras mutant MEFs. Texture analysis of MEF cells labeled with DAN13aa and visualized in TIRF. Data are shown as average intensity versus texture value (a.u.) per cell. Data from 2 independent experiments.

## Discussion

Establishment of PIP heterogeneity is a hallmark of several cellular processes. The spatial heterogeneity spans several orders of magnitude from signaling and trafficking events to cell-wide polarization (1–3, 35, 36). The extent to which cellular signaling programs is sensitive to heterogeneities in local PI(4,5)P_2_ densities has been speculated but remains incompletely explored. It is possible that clustering of PI(4,5)P_2_ enhances some signaling activities or that high densities of some proteins, especially those that are membrane anchored, could confine the lipids themselves (12, 19). Membrane PI(4,5)P_2_ hotspots could, alternately, be a site for conversion to PI(3,4,5)P_3_ as expected in Ras-driven membrane recruitment and activation of PI3K.

We sought to apply a recently introduced peptide-based sensor for PI(4,5)P_2_ detection to quantification of PI(4,5)P_2_ densities. Using a reconstituted, stacked supported lipid bilayer system, we demonstrated that the sensor is sensitively responsive to very low, physiologically-relevant levels of PI(4,5)P_2_ and that the ratiometric signal scales linearly with the lipid density. The fast binding kinetics should be minimally perturbative to PI(4,5)P_2_-dependent processes. Further, the establishment of a calibration curve is bolstered by the facile application of the sensor to living cells given the high membrane permeability.

Our observation of PI(4,5)P_2_ heterogeneity is similar to reports of enrichment in PI(3,4)P_2_ in response to PDGF stimulation and may hint at a broader design principle regulating PIP dynamics ((37). When considering specific Ras mutations in an isogenic line, we found a surprising clustering of PI(4,5)P_2_ spatial distributions and mean PI(4,5)P_2_ levels. High PI(4,5)P_2_ densities in G13D mutant cells as compared to G12 mutants may suggest enhanced PTEN-based conversion of PI(3,4,5)P_3_ to PI(4,5)P_2_. This agrees with data showing that elevated PI3K activity is opposed by PTEN phosphatase (37). The levels and heterogeneity of mutant cells is notably shifted from wild-type cells, which may indicate that changes in PI(4,5)P_2_ regulation at the membrane is a hallmark of Ras mutant cells.

Increased PI(4,5)P_2_ density is expected may enhance many signaling pathways, in addition to serving as a substrate for conversion to PI(3,4,5)P_3_ as discussed above. How these competing programs are affected by spatial heterogeneity in PI(4,5)P_2_ is unknown. We speculate that this may be a key source of regulating and partitioning cellular activities. Local increases in PI(4,5)P_2_ density would be expected to drive activation of signaling from membrane condensates such as LAT assemblies in T cells and proposed EGFR assemblies in epithelial cells through SOS activation and priming MAPK signaling (7, 38, 39). Whether PI(4,5)P_2_ is sequestered within these assemblies or can be hydrolyzed to drive disassembly, the role for these rare lipids may be highly consequential.

## Materials and Methods

### DAN13aa sensor

The DAN13aa sensor was produced and purified according to previously established methods (20).

### Cell Culture

All cells were maintained at 37°C and 5% CO_2_. Isogenic mouse embryonic fibroblasts (MEFs) were cultured as described previously (21). Briefly, cells were cultured in phenol-free, high glucose Dulbecco’s Modified Eagle Medium (DMEM) supplemented with 10% (v/v) fetal bovine serum (FBS), 100 IU/mL penicillin, 100 μg/mL streptomycin, 2 mM L-glutamine, and 1 mM sodium pyruvate.

### Preparation of Small Unilamellar Vesicles (SUVs) and Imaging Chamber Assembly

On the day of experiments small unilamellar vesicles (SUVs) with the following composition were formed via tip sonication using lipids obtained from Avanti Polar Lipids (Alabaster, AL).

| Lipid Composition of Small Unilamellar Vesicles |  |  |  |
| --- | --- | --- | --- |
| Sample | 1,2-dioleoyl-sn-glycero-3-phosphocholine (DOPC) | 1,2-dioleoyl-sn-glycero-3-phospho-L-serine (DOPS) | L-α-phosphatidylinositol-4,5-bisphosphate (PIP<sub>2</sub>) |
| 0% PIP<sub>2</sub> | 99 mol% | 1 mol% | 0 mol% |
| 0.5% PIP<sub>2</sub> | 99.5 mol% | 0 mol% | 0.5 mol% |
| 1% PIP<sub>2</sub> | 99 mol% | 0 mol% | 1 mol% |
| 4% PIP<sub>2</sub> | 96 mol% | 0 mol% | 4 mol% |
| 10% PIP<sub>2</sub> | 90 mol% | 0 mol% | 10 mol% |

Following sonication, the lipids were centrifuged for 30 – 60 minutes at ∼23,000 x g at 4°C. The resulting supernatant was then transferred to a fresh tube to avoid potential contamination from the titanium pellet. Lipids were stored on ice until needed.

25 mm #1.5 coverslips (Warner Instruments, Holliston, MA) were sonicated for 30 minutes in 50:50 isopropyl alcohol:ddH_2_O, and then rinsed thoroughly with ddH_2_O. The coverslips were then etched for 5 minutes with freshly prepared piranha solution (3:1 H_2_SO_4_:H_2_O_2_) and rinsed thoroughly with ddH_2_O. The coverslips were then dried with N_2_ and assembled into an Attofluor™ Cell Chamber (Invitrogen, Waltham, MA).

### Stacked Supported Lipid Bilayers (SSLB)

Construction of stacked supported lipid bilayers (SSLB) was adapted from (22) and following the schematic shown in Figure 1C. Assembled imaging chambers were coated with 0.1 mg/mL poly-D-lysine (PDL, Gibco) and rinsed 3 × 5 mL with 1X PBS. This was followed by one final rinse with 10X PBS. 750 mL was removed and 250 mL of freshly prepared 0 mol% PIP_2_ SUVs in aqueous suspension was added to the chamber resulting in a final lipid concentration of 0.5 mg/mL. Using a Pasteur pipette, one drop (∼20 uL) of 6M HCl was added to each chamber and incubated at room temperature for 4 minutes. The increased salt concentration and lower pH was shown to be required to drive vesicle fusion and form a contiguous bilayer (data not shown). The chamber was then rinsed 3 × 5 mL 1X PBS. The next layer of PDL was diluted 1:10 into the chamber prior to rinsing with 1X and 10X PBS as before. The final bilayer (0 – 10% PI(4,5)P_2_) was then deposited as described above. The resulting stacked bilayers were imaged immediately.

### Characterization of Stacked Supported Lipid Bilayers (SSLB)

Construction of stacked supported lipid bilayers (SSLBs) according to Figure 1C was validated in two separate dual-color experiments, one for the lipid bilayers and another for the poly-lysine layers. For these experiments, 0% PIP2 SUVs were used for both lipid bilayers. After deposition, the first lipid bilayer was stained with the green lipophilic carbocyanine dye, DiO (λ_Ex_/λ_Em_ = 484/501 nm, ThermoFisher Scientific). The sample was then imaged in TIRF on piranha-etched coverslips. Fluorescence recovery after photobleaching (FRAP) was conducted to confirm mobility of the bilayer (Figure SX). The same process was repeated in the same sample for the top bilayer using the far-red fluorophore, DiD (λ_Ex_/λ_Em_ = 644/663 nm, ThermoFisher Scientific).

For the poly-lysine layers, a 25% (m/m) solution was created by doping either poly-L-lysine-Cy3 (λ_Ex_/λ_Em_ = 554/566, NanoCS) or poly-L-lysine-Cy5 (λ_Ex_/λ_Em_ = 647/665, NanoCS), for the bottom layer or top layer, respectively, into non-fluorescent poly-D-lysine (Gibco). The sample was then imaged in TIRF on piranha-etched coverslips. FRAP was conducted on each layer (Figure SX).

### Calibration of DAN13aa in Stacked Supported Lipid Bilayers (SSLB)

Freshly prepared stacked supported lipid bilayers (SSLBs) with 0 – 10 mol% porcine brain PI(4,5)P2 in the top bilayer were treated with 10 nM (final concentration) of the PI(4,5)P2 peptide-based biosensor, DAN13aa, and incubated at room temperature for 10 – 15 minutes. Samples were then mounted on the Nikon Eclipse Ti2 microscope fitted with a room temperature stage and a 100X oil TIRF objective (numerical aperture = 1.49). Each sample was excited using a 405 nm laser in TIRF and emission was collected at 405 nm (405-405, PIP2-bound) and 488 nm (405-488, unbound). All data were presented as a ratio of I_405-405_ to I_405-488_ to represent the bound fraction of the peptide. Samples were exposed for 50 ms at 10 mW. Samples were then washed 3 × 5 mL with 1X PBS and imaged again using identical settings in a new field of view (FOV). Pre-wash data was used to construct the calibration curve presented above.

### Immunofluorescence

Mouse embryonic fibroblasts (MEF) G12D and WT cells were plated onto glass coverslips pre-coated with 0.1% poly-L-lysine solution to increase cell adherence. The cells were washed three times with 1x Phosphate Buffer Saline (PBS). The cells were then fixed at room temperature for one minute using 0.5% Paraformaldehyde (PFA) solution in PBS. Afterward, the cells were washed three times with 1x PBS. Following fixation, cells were permeabilized for three minutes using 0.2% Triton X-100 solution in PBS at room temperature to facilitate penetration of antibodies into the cells. The cells were washed three times with 1x PBS. A 1% m/v solution of Bovine Serum Albumin(BSA) in PBS was prepared and was used to block cells for 15 minutes to prevent non-specific antigen binding. The Mouse Anti-PI(4,5)P_2_ (primary antibody) was diluted to 10 ug/mL into BSA solution. The primary antibody was incubated for 30 minutes at room temperature. Cells were then subjected to three washes with 1x PBS. The goat anti-mouse A555(Secondary Antibody) was diluted to 10 ug/mL in the BSA solution. The secondary antibodies were incubated onto the cells at room temperature for one hour. Cells were washed again three times with 1x PBS. To visualize the PIP2 antigen localization, total internal reflection fluorescence TIRF microscopy was used. Specifically, the Epi 560 channel was utilized to visualize Yap fluorescence on the secondary antibodies.

### *In cellulo* Measurements of PIP2 Using DAN13aa

Isogenic mouse embryonic fibroblasts (MEFs) were plated on sterilized, poly-D-lysine coated, #1.5 glass coverslips in complete, phenol-free media 12 – 18 hours prior to imaging. Coverslips were then assembled into an Attofluor™ Cell Chamber (Invitrogen, Waltham, MA) and mounted on the Nikon Eclipse Ti2 microscope fitted with an incubated stage (Tokai Hit; humidified, 37°C + 5% CO_2_) and a 100X oil TIRF objective (numerical aperture = 1.49). The DAN13aa sensor was added to chamber at 1 µM final concentration and allowed to incubate for 15 minutes. Each sample was excited using a 405 nm laser in TIRF and emission was collected at 405 nm (405-405, PIP2-bound) and 488 nm (405-488, unbound). Cells were also imaged using the label-free imaging modality, reflection interference contrast microscopy (RICM).

### Phosphate Quantification Assay

Assay was modified from previous reports. Clean glassware with 50% H_2_SO_4_ solution. Prepare Ammonium Molybdate/Malachite Green working solution (3:1 by volume 0.2% Malachite Green in MQ H_2_O:4.2% Ammonium Molybdate in 5 M HCl). Stir for 30 minutes, then filter with Whatman #2 Filter Paper and store in the dark. The working stock is stable for approximately 1 week. Prepare 0, 1, or 4 mol% PIP_2_-containing SUV samples following procedure above. Prepare DOPC lipid standard curve samples from 1-10 nmols and dilute with working solvent (3 CHCl_3_: 2 MeOH). Aliquot samples to test tubes in triplicate. Evaporate off solvent in Cole-Parmer BH-200 Series Stuart Dry Block Heater set to 180°C. Sample preparation was optimized for each PIP_2_-containing sample. Let cool, then add 50 mL of concentrated perchloric acid to samples, 80 mL to 0 mol% SUV samples, 100 mL to 1 mol% SUV samples, and 250 mL to 4 mol% SUV samples. Heat samples for 20 min, 0 mol% SUV samples for 35 min, and 1,4 mol% samples for 45 min. Let cool. Add 400 mL of MQ H_2_O, 2 mL of Ammonium Molybdate/Malachite Green working solution, and 80 mL of 1.5% Tween 20 solution. Mix well, and measure absorbance at 660 nm.

### Microscopy

TIRF experiments were performed on a Nikon Eclipse Ti2 inverted microscope equipped with a motorized Epi/TIRF illuminator, Perfect Focus system, and a motorized stage. Imaging was performed using a 100X oil TIRF objective (numerical aperture = 1.49). Multicolor imaging used a quad-color excitation cube. A reflection interference contrast (RICM) excitation cube was used for high contrast characterization of substrates, and cell membranes. Images were captured on an EM-CCD (iXon Lite L897, Andor Inc.). All samples treated with 10 nM DAN13aa, excited using a 405 nm laser in a TIRF configuration and emission was collected at 405 (405-405) and 488 (405-488) nm. Exposure times, multidimensional acquisitions, and time-lapse periods were set using Nikon Elements software.

### Data Analysis

All analysis was performed using functions in MATLAB equipped with the image processing toolbox DipImage. Images were background and flat-field corrected prior. For images of bilayers, data acquired as image tiles were cropped to single fields of view. Ratiometric analysis was carried out by image division in a pixelwise manner. Calibrations were based on mean intensities across at least 25 fields of view in at least 3 independent experiments using unique stacked supported lipid bilayers.

For cell data, cells were masked using the reflection interference contrast (RICM) image. Masked data were used in ratiometric images.

For texture analyses, average pixel intensity in the cell mask was calculated and plotted against the gray-level co-occurrence matrix (GLCM). The GLCM contrast was implemented in MATLAB.

The formula for GLCM is:

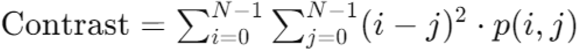

