## Supplemental Figures for "Direct measurement of PIP_2_ densities in biological membranes using a peptide-based sensor"

**Supplemental Information**

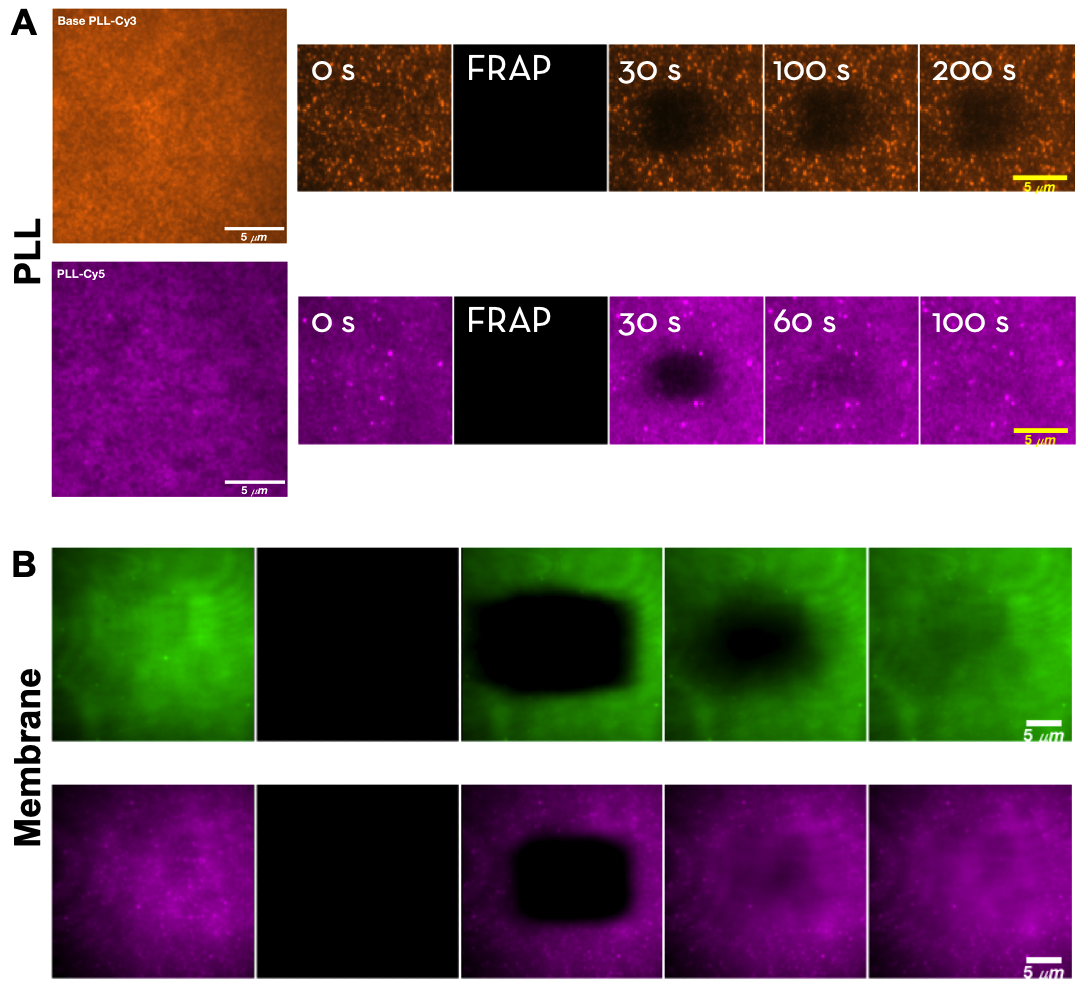

**Supplemental Figure 2**. Mobility of components of stacked supported bilayer configuration. Characterization of mobility of the components of SSLBs by fluorescence recovery after photobleaching (FRAP). A. Fluorescent poly-D-lysine (PDL-Cy3) on etched glass was immobile (top), whereas the PDL-Cy5 between the two SLBs was mobile.

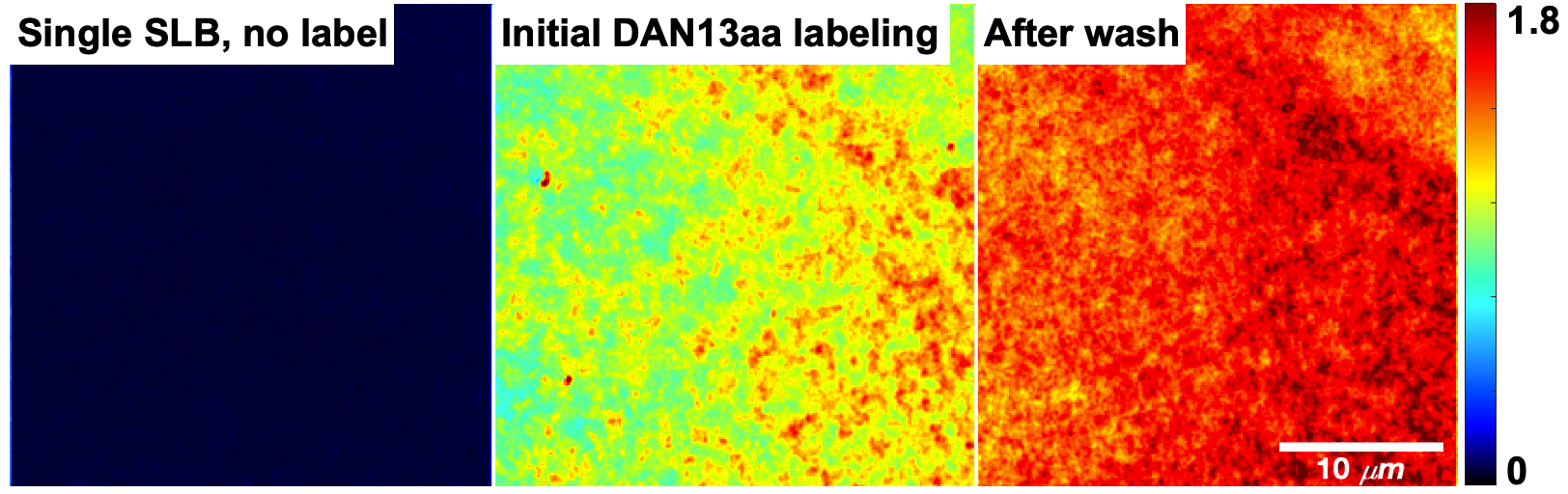

**Supplemental Figure 1**. DAN13aa sensor accumulation on glass displaying a single SLB. Representative, ratiometric snapshots of a single supported lipid bilayer on etched glass before, during initial exposure to DAN13aa, and after washing, following 10 minutes of incubation with the sensor, respectively.

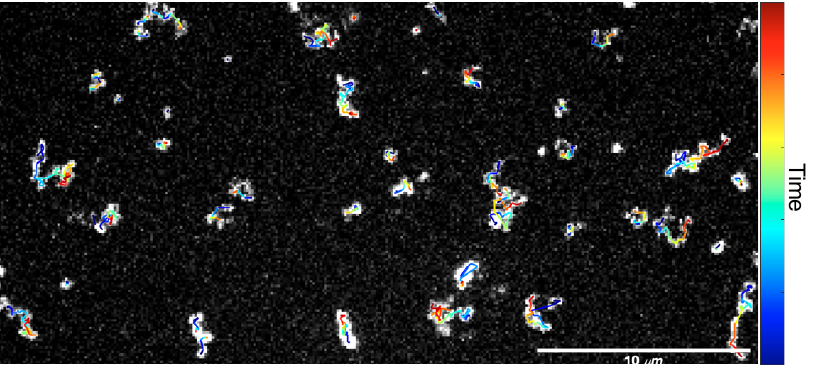

**Supplemental Figure 3**. Single PIP2 diffusion in a stacked SLB configuration. PIP2-TopFluor was doped into the top SLB in the stacked configuration at 4 mole percent. Following bleaching down to single molecule levels, individual lipids could tracked.

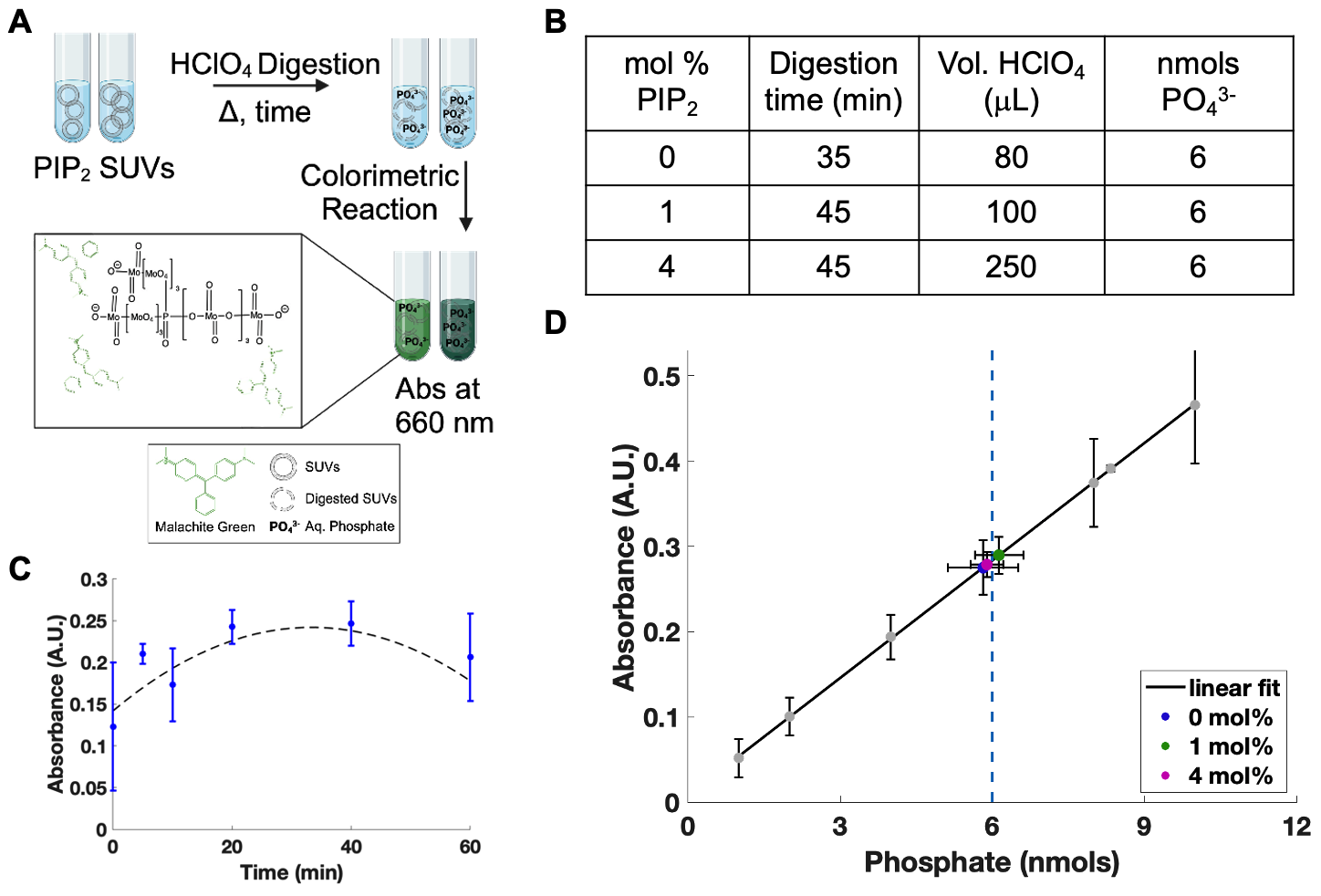

**Supplemental Figure 4**. Verification of reconstituted PI(4,5)P_2_ content in membranes using a colorimetric assay. A. Schematic of assay. Vesicles were prepared and digested with perchloric acid prior to application of the colorimetric Malachite Green reagent. B. Table of vesicle compositions and digestion times. C. Digestion times were optimized to achieve peak signal detection. D. Measurement of phosphate amounts in samples using assay. Each sample targeted a final amount of phosphate to be 6 nanomoles.

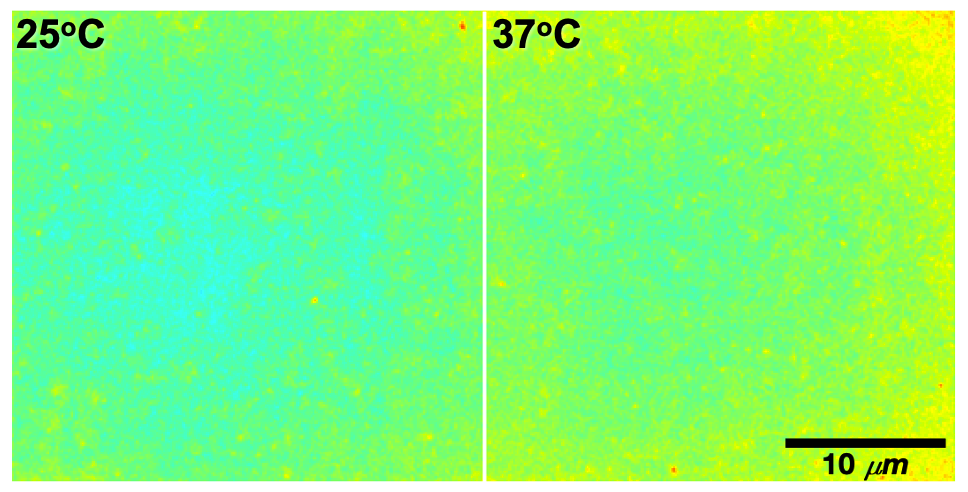

**Supplemental Figure 5**. DAN13aa sensor performance as a function of temperature. Ratiometric images for stacked SLBs imaged at either 25^o^C or 37^o^C.

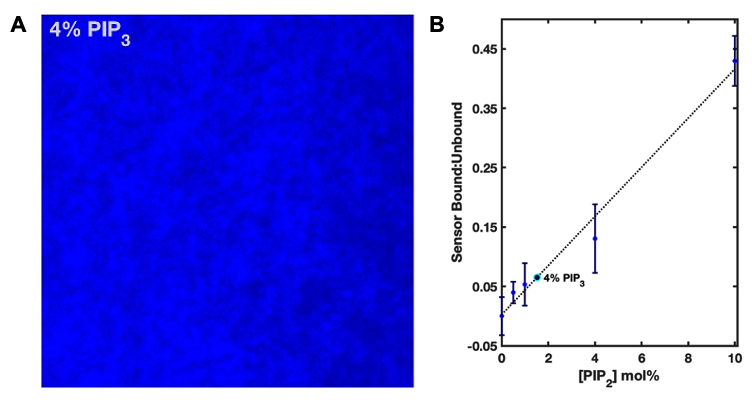

**Supplemental Figure 6**. DAN13aa sensor detection of PIP3. A. Image of stacked supported lipid bilayer containing 4 mol % PIP_3_. B. Ratiometric signal from DAN13aa sensor due to PIP3 plotted on the standard curve.

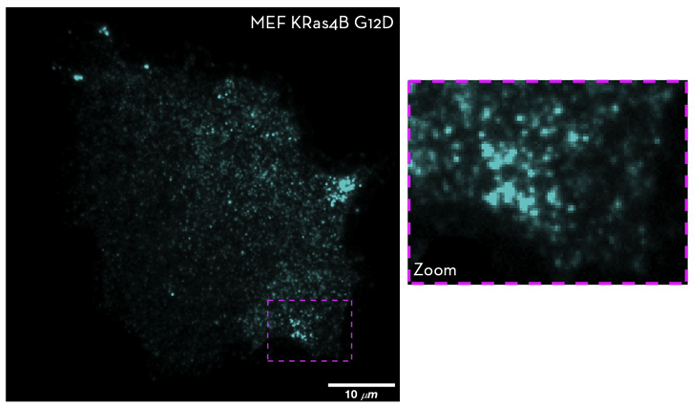

**Supplemental Figure 7**. IF and heterogeneity in mutant membranes. Representative images of MEF KRaas4B G12D mutant cells stained for PI(4,5)P_2_ and imaged in TIRF. Zoomed region shows clustered lipids.
